# Introgressions lead to reference bias in wheat RNA-Seq analysis

**DOI:** 10.1101/2023.10.04.560829

**Authors:** Benedict Coombes, Thomas Lux, Eduard Akhunov, Anthony Hall

**Affiliations:** Earlham Institute, Norwich, Norfolk, NR4 7UZ, UK; Plant Genome and Systems Biology, Helmholtz Zentrum München, Neuherberg, Germany; Department of Plant Pathology, Kansas State University, Manhattan, KS, USA

## Abstract

RNA-Seq is a fundamental technique in genomics, yet reference bias, where transcripts derived from non-reference alleles are quantified less accurately, can undermine the accuracy of RNA-Seq quantification and thus the conclusions made downstream. Reference bias in RNA-Seq analysis has yet to be explored in complex polyploid genomes despite evidence that they are often a complex mosaic of wild relative introgressions, which introduce blocks of highly divergent genes. Here we use hexaploid wheat as a model complex polyploid, using both simulated and experimental data to show that RNA-Seq alignment in wheat suffers from widespread reference bias which is largely driven by divergent introgressed genes. This leads to underestimation of gene expression and incorrect assessment of homoeologue expression balance. By incorporating gene models from ten wheat genome assemblies into a pantranscriptome reference, we present a novel method to reduce reference bias, which can be readily scaled to capture more variation as new genome and transcriptome data becomes available.

## Background

Quantification of gene expression using RNA-Seq is a fundamental technique in genomics research. It has been employed in numerous publications across a range of biological systems to identify candidate genes underlying traits of interest, uncover transcriptional pathways and networks, and investigate hypotheses relating to gene and transcriptional evolution and adaptation. In RNA-Seq experiments, mRNA, which represents a snapshot of the expression of each gene at the time of sampling, is extracted from the biological sample, converted to cDNA, and sequenced. The number of resulting RNA-Seq reads deriving from each gene/transcript are quantified, with the number of reads proportional to the level of expression of that gene/transcript. Quantifying the expression level of each transcript and/or gene typically involves alignment of sequencing reads to the reference genome or transcriptome of the sequenced species using spliced alignment tools such as HISAT2 [1] and STAR [2] or pseudoalignment tools such as kallisto [3] and Salmon [4]. Despite these tools typically being developed and benchmarked with human data, they are widely used across numerous biological systems, often without consideration for how they will behave with specific challenges the genomes of different species present.

Making meaningful inferences from RNA-Seq data relies upon the accuracy of alignment and quantification; downstream analyses and subsequent interpretation assumes that the estimated gene expression reflects actual gene expression in the biological samples. However, nucleotide variation in the coding region of genes between the sequenced sample and the reference genome/transcriptome leads to errors in read assignment during the alignment/pseudoalignment step. Some reads may be unassigned, while others may be assigned to the wrong locus. This source of error is widely known as reference bias as transcripts derived from alleles present in the reference sequence will be quantified more accurately [5].

The impact of reference bias in RNA-Seq analysis hasn’t been assessed in complex polyploid genomes such as wheat despite these genomes having characteristics that may increase the extent and degree of reference bias relative to species with simpler genomes. Polyploidisation increases the number of alleles per gene, typically resulting in a pair of alleles, known as homoeologues, in each subgenome; however, subsequent gene duplications or deletions can change the relative copy number of homoeologues between the subgenomes. As RNA-Seq reads are derived from all subgenomes at once, read assignment must be able to distinguish reads deriving from homoeologues. Accurate discrimination of wheat homoeologue RNA-Seq reads has been demonstrated with both pseudoalignment [6, 7] (99.9% accuracy) and alignment-based (98% accuracy) [7] methods when mapping reads back to the genome from which they derived. However, when mapping reads from a different genotype, unequal divergence between homoeologues relative to the reference genome may compromise the accuracy of the expression balance estimation between homoeologues.

Introgression events, the introduction of genetic material from one species to another [8], are common among plants; in fact, its frequency is thought to be higher in plants than in animals, due to higher rates of interspecific hybridisation success [9]. Additionally, introgression is commonly used to introduce genetic variation for crop improvement [10]. Several studies have demonstrated how common introgressions are in wheat accessions with some accessions being comprised of up to 34% introgressed material [11–14]. The production of chromosome-level genome assemblies of modern elite wheat cultivars confirmed this, revealing introgressions from wild and domesticated relatives, including species outside of the *Triticum* and *Aegilops* genera, present in one or multiple cultivars [15, 16]. These introgressions introduce greater sequence divergence between varieties than observed between varieties at non-introgressed regions; this increased divergence likely leads to an increased proportion of reads that are unable to be assigned correctly.

Using simulated and experimentally-generated RNA-Seq data, we identify non-trivial levels of reference bias in RNA-Seq mapping in wheat which can largely be attributed to introgressions. This leads to incorrect estimates of relative expression between homoeologues and incorrectly called differences in expression between cultivars. By constructing a pantranscriptome reference composed of Chinese Spring transcripts and transcripts from the wheat pangenome assemblies, we demonstrate how reference bias caused by divergent alleles can be reduced.

## Results

### Reference bias in wheat is driven by divergent genes introduced via introgressions and results in underestimation of gene expression

To explore the impact of reference bias on the quantification of gene expression in wheat, we simulated 1000 read pairs from each high-confidence (HC) gene in Chinese Spring RefSeq v1.1 and the nine chromosome-level genome assemblies produced from the wheat pangenome project [15, 17] if the longest transcript of the gene is at least 500bp. These reads were pseudoaligned or aligned to the Chinese Spring reference transcriptome or genome using kallisto or STAR, respectively. These algorithms represent pseudoalignment and alignment-based methods and are among the most commonly used tools for RNA-Seq quantification in the wheat community.

Mapping Chinese Spring reads to Chinese Spring, hereafter referred to as self-mapping, yields very accurate estimates of gene expression, with kallisto slightly outperforming STAR (Figure 1a, Figure 1b, Supplementary Table 1). Using kallisto, 88401/88443 (99.95%) of genes were correctly quantified (between 500 and 1500 read pairs). 32 genes were underestimated (< 500 read pairs) and 10 genes were overestimated (> 1500 read pairs). Using STAR, 87689/88443 (99.15%) were correctly quantified with 504 and 250 genes underestimated and overestimated, respectively.

**Figure 1.**
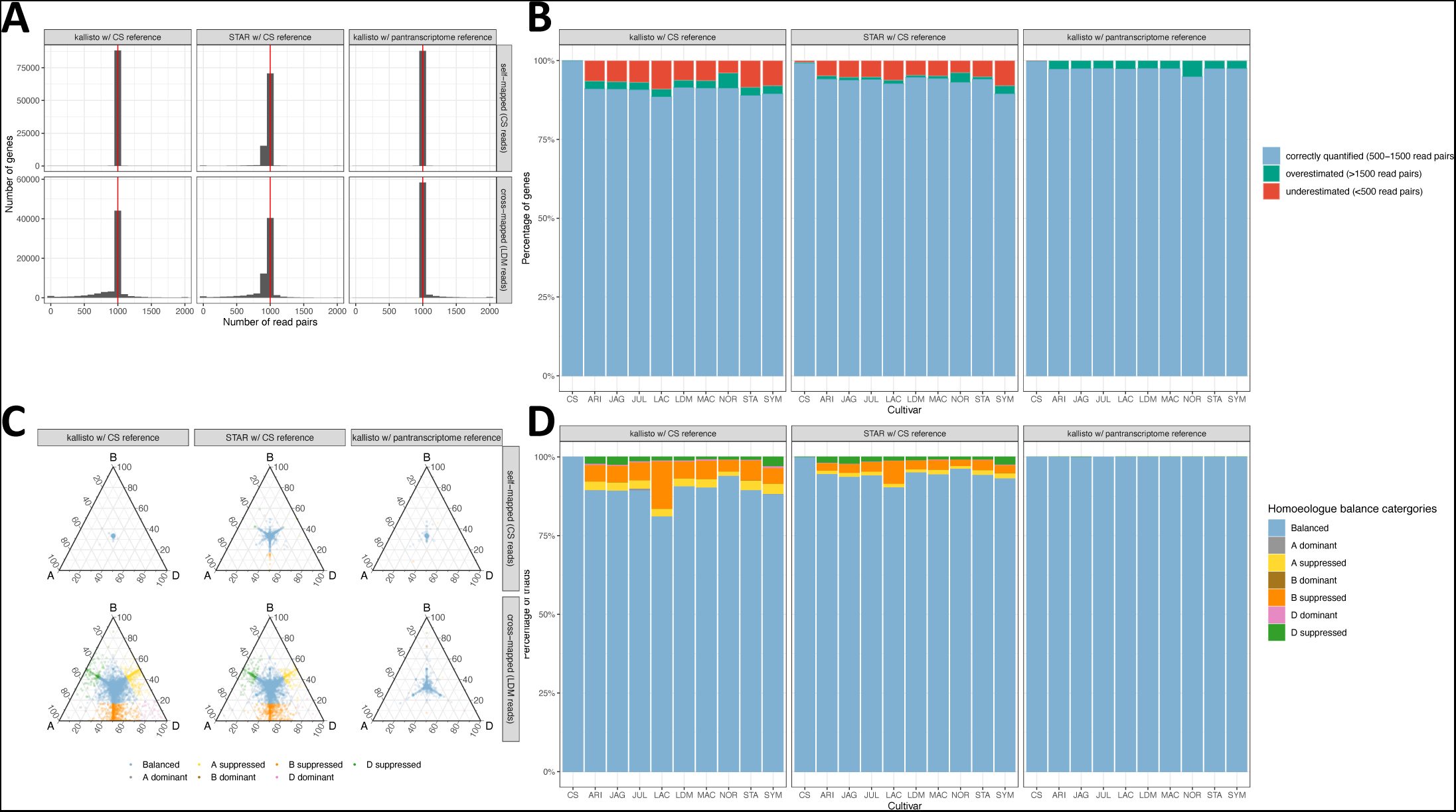
Reference bias in wheat. **A)** Distribution of read counts when self-mapping Chinese Spring simulated reads or cross-mapping Landmark simulated reads. Comparing STAR and kallisto using the Chinese Spring RefSeq v1.0 reference and RefSeq v1.1 transcriptome and kallisto using the pantranscriptome reference. **B)** Percentage of genes with expression estimated correctly, expression underestimated (< 500 read pairs) and expression overestimated (> 1500 read pairs) for simulated reads from 10 cultivars aligned to Chinese Spring with kallisto and STAR or to the pantranscriptome reference with kallisto. **C)** Balance of homoeologue expression across triads when self-mapping Chinese Spring or cross-mapping Landmark simulated reads, comparing STAR and kallisto using the Chinese Spring RefSeq v1.0 reference and RefSeq v1.1 transcriptome and kallisto using the pantranscriptome reference. Each point on the ternary plot represents one triad. Points towards a corner indicate dominant expression of that homoeologue, points opposite a corner indicate suppression of that homoeologue. **D)** Percentage of triads in each expression category, using simulated reads from 10 cultivars aligned to Chinese Spring with kallisto and STAR or to the pantranscriptome reference with kallisto.

Mapping reads generated from the other cultivars to Chinese Spring, hereafter called cross-mapping, yielded much less accurate estimation of gene expression with a skew towards underestimation (Figure 1a, Figure 1b, Supplementary Table 1). The percentage of genes correctly quantified ranged from 55773/63001 (88.53%) for Lancer, with 5700 (9.05%) and 1528 (2.43%) under and overestimated, respectively, to 58468/64077 (91.2%) for Norin61, with 2527 (3.94%) and 3082 (4.81%) genes under and overestimated, respectively. For cross-mapping, unlike self-mapping, STAR appears to perform better than kallisto; the proportion of correctly quantified genes ranged from 58390/63001 (92.68%) for Lancer, with 3916 and 695 under and overestimated, respectively, to 59648 (93.1%) for Norin61, with 2450 and 1979 genes under and overestimated, respectively.

To explore the effect of reference bias on the quantification of homoeologue expression balance, we calculated the proportion of triads belonging to each category that defines a different state of relative homoeologue expression. As reads were simulated evenly across genes, all triads should be classified as balanced; therefore, triads classified as imbalanced (one or two homoeologues with expression greater than the other(s)) are considered incorrectly classified. The percentage of correctly classified triads varies between 80.97% (Lancer) and 93.84% (Norin61) using kallisto and between 90.23% (Lancer) and 96.12% (Norin61) using STAR (Figure 1c, Figure 1d, Supplementary Table 2). Across the cultivars, triads incorrectly classified as suppressed, where one homoeologue is estimated to be expressed less than the others, were far more common than triads incorrectly classified as dominant, where one homoeologue is estimated to be expressed more highly than the others (Figure 1d, Supplementary Table 2). This reflects how the reference bias leads to more underestimated than overestimated genes.

To explore the extent of errors when comparing two cultivars mapped to a common reference, we compared the estimated expression of Lancer and Jagger genes, whose simulated reads were both aligned to Chinese Spring using STAR (Figure 2a). Genes with read counts > 1.5x or < 1/1.5x compared to the other cultivar were classified as incorrectly quantified. Using STAR, 4791/60338 (7.94%) genes were incorrectly quantified between the two cultivars; of these genes, 2747 and 2044 genes had a lower read count in Lancer and Jagger, respectively.

**Figure 2.**
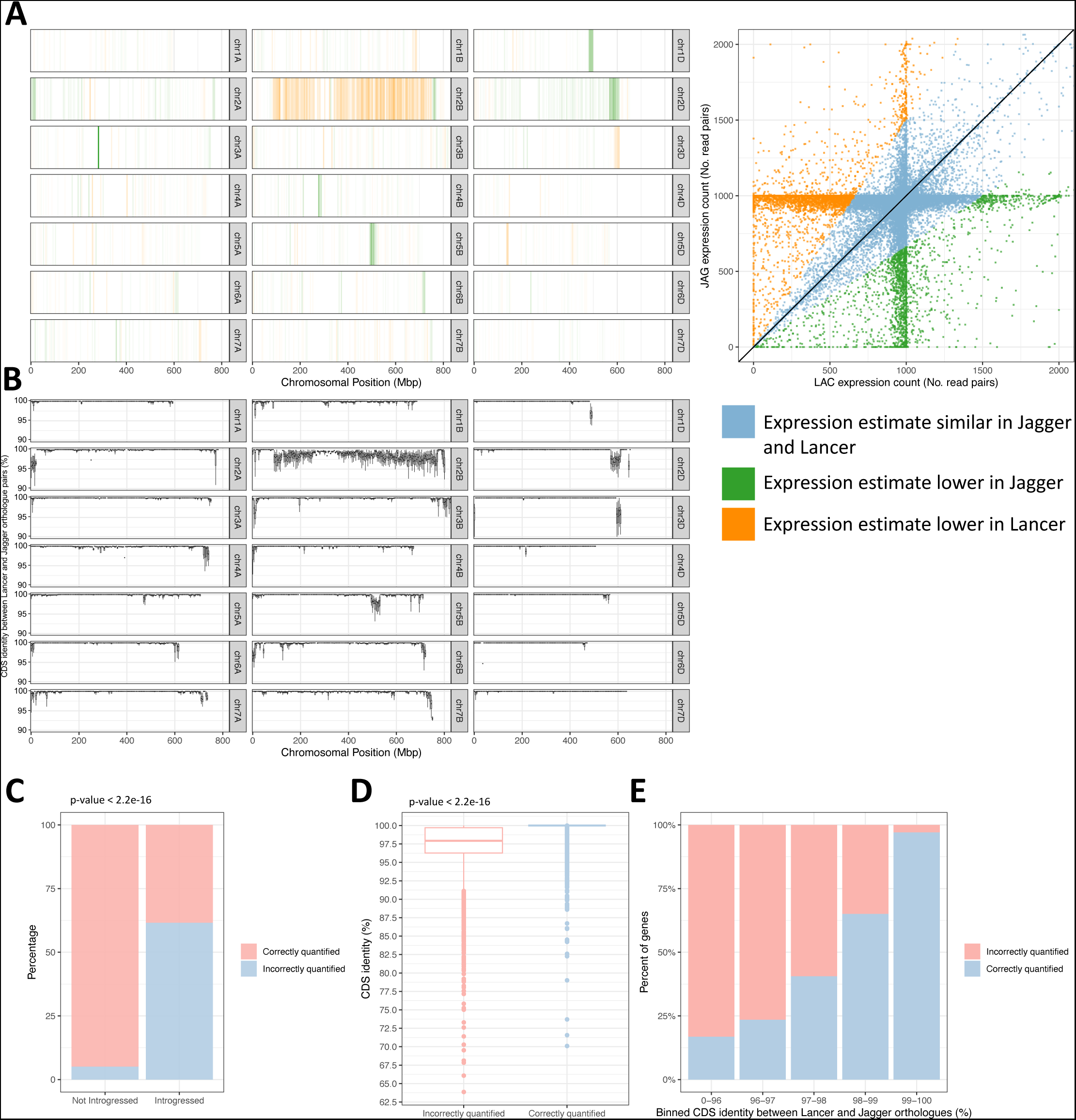
Exploring the impact of reference bias on expression differences between cultivars and enrichment of incorrectly quantified genes within introgressions. **A)** Scatter plot shows expression counts for Lancer-Jagger orthologue pairs. Genes are considered incorrectly quantified if their estimated read count is 1.5x or 1/1.5x the other cultivar. The chromosome plot shows the distribution of incorrectly quantified genes in 5Mbp windows, coloured by the cultivar in which the estimated expression is lower; orange blocks are underestimated in Lancer compared to Jagger, while green blocks are underestimated in Jagger compared to Lancer. The reads are aligned using STAR as this outperformed kallisto for cross-mapping. **B)** CDS nucleotide identity between Lancer and Jagger 1-to-1 orthologue pairs, binned into 5Mbp genomic windows based on Chinese Spring RefSeq v1.0. **C)** Percentage of genes incorrectly quantified and correctly quantified in characterised introgressed regions and regions not characterised as introgressed. **D)** CDS nucleotide identity between Lancer and Jagger 1to1 orthologue pairs for those that are incorrectly quantified and those that are correctly quantified. **E)** Percentage of genes incorrectly quantified and correctly quantified, split into bins of different levels of CDS nucleotide identity.

We observed a clear overlap between clusters of incorrectly quantified genes and regions of divergence between the cultivars (Figure 2a), identified by blocks of reduced CDS nucleotide identity between pairs of orthologues between Lancer and Jagger. Such gene-level divergence is indicative of introgressed material; indeed, several of these blocks correspond to previously characterised introgressions and likely additional introgressions that are yet to be characterised. These introgressions include (coordinates based on Chinese Spring RefSeq v1.0): *Aegilops ventricosa* introgression in Jagger (chr2A:1-24643290) [15, 18, 19]; *Triticum timopheevii* introgression in Lancer (chr2B:89506326-756157100) [15, 19]; *Aegilops comosa* introgression in Jagger (chr2D:570141481-613325841) [19]; and a *Thinopyrum ponticum* introgression in Lancer (chr3D:591971000-615552423) [15, 19]. 1173/3054 (61.59%) of introgressed genes were incorrectly quantified between the two cultivars, compared to 2910/57284 (5.24%) non-introgressed genes incorrectly quantified (chisq. p < 2.2e-16) (Figure 2c). Genes with an introgressed copy in Lancer tend to be underestimated in Lancer and genes with an introgressed copy in Jagger tend to be underestimated in Jagger.

In further support of CDS divergence being a predominant contributing factor to incorrect quantification, we found that incorrectly quantified genes have a mean CDS identity between orthologue pairs of 97.3% compared to a mean of 99.9% for genes correctly quantified (Figure 2d; p-value < 2.2e-16; 95% confidence interval ranges from 2.45 to 2.63). The percentage of genes incorrectly quantified ranges from 83.2% for genes with <96% CDS identity between orthologues to just 2.9% for genes with >=99% identity between orthologues (Figure 2e).

### Reducing reference bias by constructing a pantranscriptome reference

The wheat pangenome project generated chromosome-level de novo assembled genomes for 9 wheat cultivars in addition to the reference cultivar Chinese Spring [15]. These include numerous introgressions that are the predominant source of reference bias we observe. High-quality gene annotations for these genome assemblies have been produced [17]. We constructed a pantranscriptome reference by adding in transcripts from the pangenome cultivars to the set of Chinese Spring transcripts if the gene existed in a 1-to-1 relationship with a gene from Chinese Spring. 80211 Chinese Spring genes had at least one 1-to-1 orthologue in another cultivar, while 59639 Chinese Spring genes had a 1-to-1 orthologue in all nine other cultivars (Supplementary Figure 1) based on OrthoFinder [20] orthogroup assignments. This pantranscriptome reference was used as the transcriptome reference for kallisto pseudoalignment. After pseudoalignment, read counts and TPMs were summed across all transcripts corresponding to a given Chinese Spring gene. Kallisto splits read counts evenly across transcripts with an identical match so redundancy of transcripts does not cause problematic multi-mapping; all transcripts corresponding to a gene can thus be added.

To ensure using this pantranscriptome reference doesn’t introduce any additional mapping errors from adding redundant transcripts, we compared quantified expression counts between four difference references: Chinese Spring, the pantranscriptome reference, Chinese Spring plus Landmark, and the pantranscriptome reference minus Landmark. The simulated reads from Landmark were used for pseudoalignment. The pantranscriptome reference performed the best, with 97.53% of genes correctly quantified. Chinese Spring plus Landmark transcripts was very similar, with 97.50% of genes correctly quantified. This demonstrates that adding redundant transcripts and summing the read counts doesn’t introduce errors in the kallisto mapping. Using the pantranscriptome reference without Landmark transcripts resulted in slightly lower level of correct quantification, with 96.84% correctly quantified. The difference is likely due to uniquely introgressed genes in Landmark that are not present in the other cultivars. However, due to many introgressed genes being common between cultivars, it still performed much better than just using Chinese Spring, which had 91.43% genes correctly quantified.

Using the pantranscriptome reference instead of Chinese Spring to quantify expression from the simulated RNA-Seq reads resulted in much more accurate quantification for genes that were previously underestimated when cross-mapping, removing nearly all gene counts below 1000 (Figure 1a, Figure 1b). There was little change in the number of genes overquantified when cross-mapping and little difference in the distribution of read counts when self-mapping (Figure 1a, Figure 1b). The distribution of read counts shows that for Lancer, the most error prone cultivar, the number of genes correctly quantified increased from 58390/63001 (92.68%) using STAR to 61352/63001 (97.38%) using the pantranscriptome reference. Using the pantranscriptome reference, only 2 genes remained quantified below 500 read pairs compared to 3916 genes when using the Chinese Spring reference. The number of triads correctly assigned to the balanced expression category also greatly increased when using the pantranscriptome reference (Figure 1d). All cross-mapped cultivars had at least 99.89% triads correctly assigned as balanced; this compares to between 80.97% and 93.84% using kallisto, and between 90.23% to 96.12% using STAR to align to Chinese Spring.

Comparing Jagger and Lancer as before, this approach reduced the number of genes incorrectly quantified in one cultivar from 5049/60338 (8.96%) to 617 (1.02%) (Supplementary Figure 2). Only 23 genes (0.0381%) remain incorrectly quantified due to underestimation in one cultivar compared to 4882 (8.90%) when aligned to Chinese Spring using STAR. Almost all the remaining error in both cross-mapped read counts and incorrectly quantified genes between cultivars is due to overestimation of gene expression, likely caused by copy number variation or presence/absence variation between cultivars, as opposed to divergence between orthologous gene models.

### Exploring reference bias caused by introgressions in experimentally-generated RNA-Seq data

Simulated RNA-Seq data is unlikely to capture the complete picture of a real experiment [21]. While our simulations highlight theoretical errors, it is important to assess how reference bias impacts published findings and how using the pantranscriptome reference corrects errors in real data. We reanalysed the data from He *et al.* (2022), in which RNA-Seq data from 198 diverse wheat accessions, alongside enrichment capture paired-end DNA reads was used to uncover eQTLs linked with homoeologue expression bias and variation in important productivity traits. Crucially for our work, they identified a set of genes whose expression exhibited negative correlation with its homoeologue across the panel. A subset of accessions possessed lowly expressed alleles in one of the homoeologues and the presence of the lowly expressed alleles was linked to various important productivity traits. This set contains 59 genes to which we have added *ELF3-D1*. While *ELF3-D1* didn’t fall into the set of very negatively correlated 59 genes, it was used as case example due to its agronomic significance. Also, it still did show a negative correlation with its B homoeologue, with this expression bias associated with agronomic traits. This set of 60 genes is hereafter referred to as genes showing lack of expression correlation.

Firstly, to identify potential introgressed regions within these accessions, we mapped the enrichment capture paired-end DNA reads to Chinese Spring RefSeq v1.0 and for each 1Mbp genomic window, calculated the mapping coverage deviation between each line and the median for that window across the accessions. Windows with a coverage deviation value significantly below 1 in an accession were labelled as possessing an introgression or deletion. The genes showing lack of expression correlation identified by He et al. (2022) are enriched in genomic windows identified introgressed or deleted (Figure 3b), with 78.2% of these genes in a genomic window identified as introgressed or deleted in 30 or more accessions. This compares to the rest of the genes in the genome, of which only 12.3% are found in a genomic window identified as introgressed or deleted in 30 or more accessions.

**Figure 3.**
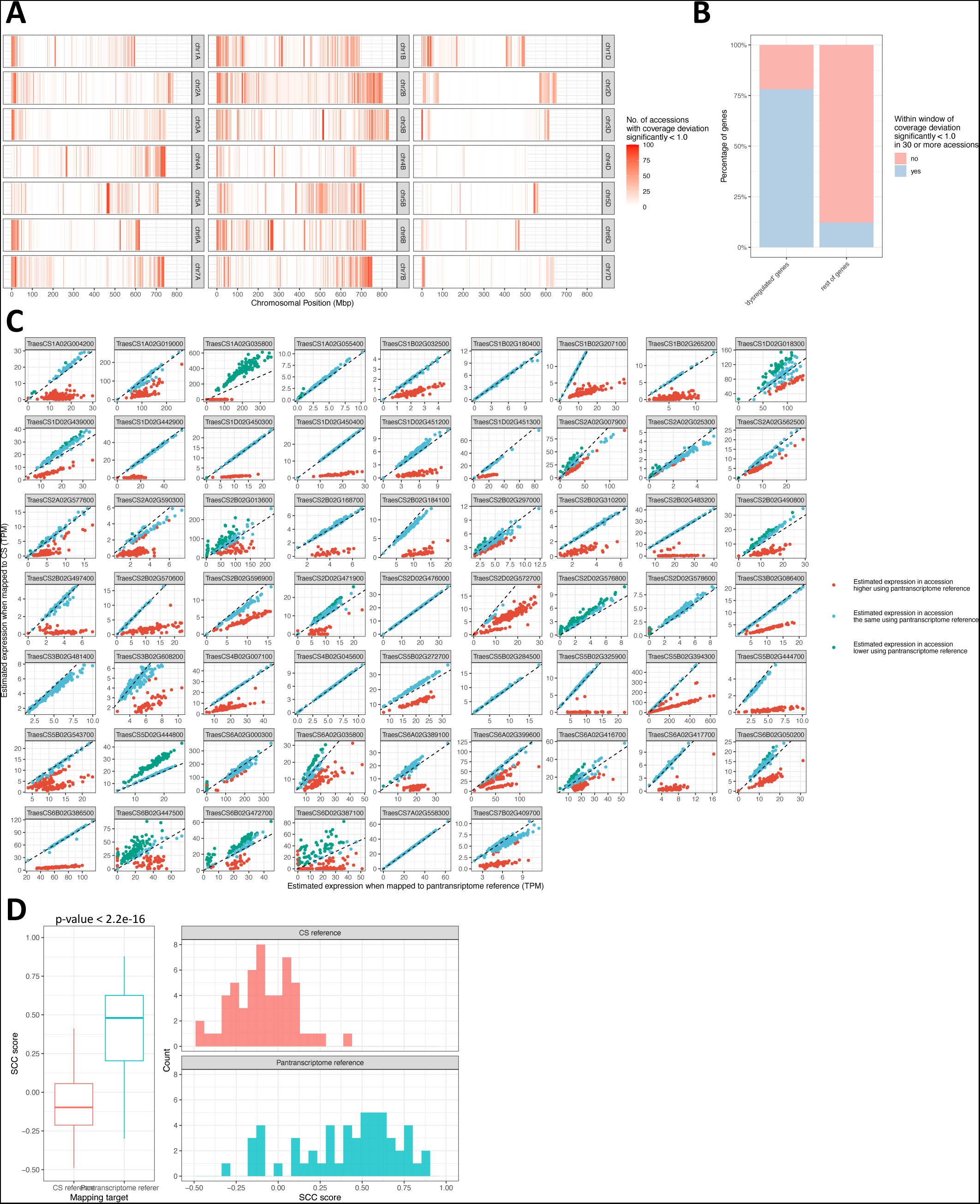
Exploring the impact of reference bias on experimental RNA-Seq data. **A)** Chromosomal distribution of the number of lines in each 1Mbp genomic window which had mapping coverage deviation significantly less than 1. **B)** The number of genes from the set of 60 genes showing lack of expression correlation identified by He et al. (2022) that are present in genomic windows identified as introgressed or deleted in 30 or more accessions. **C)** Estimated expression of the 60 genes showing lack of expression correlation identified by He et al. (2022), using either the Chinese Spring RefSeq v1.1 transcriptome or the pantranscriptome reference as targets for kallisto pseudoalignment. The dashed black line represents x=y, which is the expected value if the reference is not affecting the estimation of gene expression. Red dots and green dots represent accessions in which a given gene has a TPM value <50% or >150%, respectively, when mapping to Chinese Spring than when mapping to the pantranscriptome reference. **D)** Spearman’s correlation coefficient between homoeologue pairs where one is the set of genes showing lack of expression correlation identified by He et al. (2022). SCC scores were computed between AB, AD and BD homoeologue pairs and the lowest score used. Triads in which any were not present in the RefSeq v1.0 HC gene annotation were excluded. The significance of the difference between SCC scores when using the Chinese Spring reference compared to when using the pantranscriptome reference was calculated using a two-tailed t-test with no assumption of equal variance.

To explore the impact of the pantranscriptome reference on estimated expression, we pseudoaligned the leaf RNA-Seq data from the 198 wheat accessions to both Chinese Spring and to the pantranscriptome reference. Kallisto was used for aligning to Chinese Spring instead of STAR for consistency with (He *et al.*, 2022). 42/60 (70%) of genes showing lack of expression correlation (fig 3c) have, in 25 or more accessions, an estimated expression less than half when mapping to Chinese Spring compared to when mapping to the pantranscriptome reference (Figure 3b). These are likely introgressed genes whose expression is underestimated when using Chinese Spring as the reference. 7/60 (11.7%) of the genes have, in 25 or more accessions, an estimated expression more than double when mapping to Chinese Spring compared to when mapping to the pantranscriptome reference (Figure 3c).

While this shows that using Chinese Spring as the reference leads to underestimation of many of these genes, it is important to look at the impact of this on the calculated correlation between homoeologues that led to them being classified as genes of interest by He et al. (2022). We found that the SCC score between homoeologues from this set was - 0.0990 when using the Chinese Spring reference and 0.407 using the pantranscriptome reference (Figure 3d; p-value < 2.2e-16; 95% confidence interval ranges from -0.603 to - 0.410). Even though this SCC value remains lower than the mean SCC (∼0.8) reported for the entire set of homoeologues (He et al., 2022), it indicates that the usage of pantranscriptome as reference increases expression correlation estimates between homoeologues compared to single reference estimates.

Several regions of coverage deviation overlap precisely with characterised introgressions from cultivars in the wheat pangenome. One such introgression is at the end of chr1D (484,302,410bp-495,453,186bp, based on RefSeq v1.0 coordinates), present unbroken in 53/198 (26.8%) accessions (Supplementary Table 3) and shared with cultivars Jagger and Cadenza from the wheat pangenome (Figure 4a). We looked at this introgression as its size is invariable between accessions. Additionally, this region was highlighted in He et al. (2022) as it contains 6 of the genes showing lack of expression correlation, including *ELF3-D1*, which was used as a case example due to its role in heading date [22]. (He *et al.*, 2022) suggest this is a terminal deletion; however, [23], identified that the terminal region, including *ELF3-D1*, is introgressed in Cadenza and Jagger, deriving from either *Triticum timopheevii* or *Aegilops speltoides*, based on the *ELF3-D1* gene model possessing an intronic deletion shared with both of these species. We can exclude *Ae. speltoides* as the donor species as protein alignments between the Jagger introgression and *Ae. speltoides* proteins showed a median protein identity of just 91.6%. As *T. timopheevii* does not have a genome assembly available, we can’t confirm it is the donor; however, the mapping profile of *T. timopheevii* reads to the Jagger genome assembly suggest it is a likely match (Supplementary Figure 3). As we can’t be certain about the donor species, we will hereafter refer to this introgression as the chr1D introgression.

**Figure 4.**
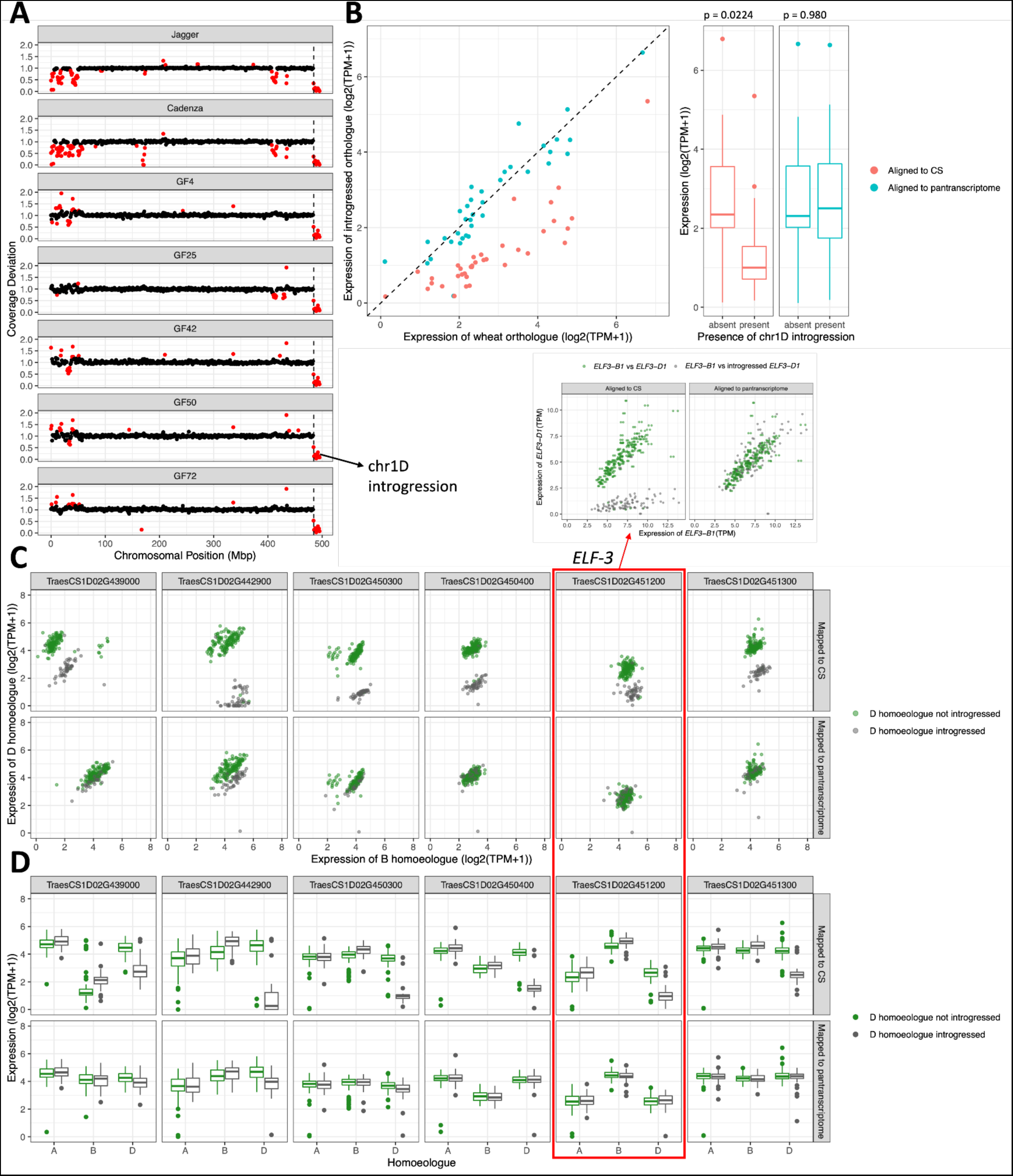
Introgressed genes falsely identified as less expressed due to reference bias. **A)** Mapping coverage deviation across chr1D of Jagger, Cadenza, and 5 accessions from He et al. (2022). The coverage drop at the end of chr1D, the left-hand border of which is indicated by the vertical dashed black line, is an introgression common to 53 of the 198 accessions and Jagger and Cadenza from the wheat pangenome. **B)** Expression of the wheat gene compared to its introgressed orthologue from the chr1D introgression, using either Chinese Spring or the pantranscriptome reference as targets for kallisto pseudoalignment. Orthologue pairs with TPM < 1 in both the introgressed and the wheat copy when mapping to the pantranscriptome reference were excluded. The significance of the difference between introgressed and non-introgressed orthologues when using the Chinese Spring or the pantranscriptome reference was calculated using two-tailed t tests with no assumption of equal variance **C)** Estimated expression level of introgressed D homoeologues compared to the wheat B homoeologues and wheat D homoeologues compared to wheat B homoeologues, using either Chinese Spring or pantranscriptome reference as targets for kallisto pseudoalignment. **D)** Expression level of triads from where the D homoeologue is an introgressed gene in a subset of lines, using either Chinese Spring or the pantranscriptome reference as targets for kallisto pseudoalignment.

We compared the mean expression of genes from the chr1D introgression across accessions that possess the introgression to their 1-to-1 wheat orthologue across the accessions lacking the introgression. When using the Chinese Spring reference, the introgressed genes appear to be less expressed than their wheat orthologues; however, when using the pantranscriptome reference, no significant difference in expression was found between the genes (Figure 4b, Supplementary Table 4).

Earlier, using simulated data, we demonstrated that reference bias can lead to incorrect assignment of expression balance across triads. To examine this phenomenon in real data, we examined the estimated expression across triads within the chr1D introgression that are also in the set of genes showing lack of expression correlation identified by He et al. (2022). When the RNA-Seq reads are pseudoaligned to Chinese Spring, in lines with the chr1D introgression, *ELF3-D1* appears to be lowly expressed and the expression of *ELF3-B1* appears slightly elevated compared to accessions without the chr1D introgression. However, when mapped to the pantranscriptome reference, the expression of *ELF3-D1* and *ELF3-B1* in accessions with the chr1D introgression appears very similar to that in accessions without the chr1D introgression (Figure 4c, Figure 4d). The CDS sequence for *ELF3-D1* from the introgression shares 97% sequence identity with *ELF3-D1* in Chinese Spring, 97.56% identity with *ELF3-A1* and 97.8% identity with *ELF3-B1*. The high divergence of *ELF3-D1* from the introgression and *ELF3-D1* from Chinese Spring and the greater similarity between *ELF3-D1* from the introgression with *ELF3-B1* from Chinese Spring explains how most reads were unable to be assigned, yet some were incorrectly assigned to the *ELF3-B1*, hence the slight increase in estimated expression of *ELF3-B1* when using the Chinese Spring reference. The five other genes showing lack of expression correlation within the chr1D introgression also showed reduced homoeologue imbalance using the pantranscriptome reference and expression level in line with accessions without the chr1D introgression, in which the triad does not contain an introgressed D homoeologue. Four of these genes also showed a slight decrease in estimated expression in the B homoeologue when mapping to the pantranscriptome reference, supporting the idea that false mapping from the introgressed gene to its homoeologue will be driving false negative correlation scores in addition to artificially low expression of the introgressed homoeologue.

## Discussion

In the emerging era of plant pangenomics, chromosome-level assemblies are being generated for an increasing number of cultivars/accessions, which will facilitate a shift away from reference genome-centric methods. Here we have demonstrated the importance of utilising these resources effectively for RNA-Seq analyses in wheat to reduce reference bias.

### RNA-Seq reference bias in wheat

Quantification of gene expression from RNA-Seq reads in wheat is very accurate when the matching reference genome for the sample is available. However, cross-mapping RNA-Seq reads leads to detectable levels of reference bias. This was seen both at the individual gene level and also when assigning triads to categories of homoeologue expression balance. A major cause of this bias appears to be introgressions of diverged haplotypes of genes from wheat’s wild and domesticated relatives. In some cases, references bias within introgressions could be severe enough to have a strong impact on downstream analyses and conclusion drawn based on these analyses. This analysis was conducted on wheat but other species with substantial introgressed content and/or polyploid genomes may suffer from the same problem. Similar analyses on other species may thus provide value for their respective communities.

Kallisto performed better for self-mapping but when cross-mapping, STAR was better able to deal with divergence between genes, although was far from resolving the issue of reference bias. Similar limitations of alignment-free methods have been previously discussed; for example, [24] demonstrated that kallisto performs poorly for lowly expressed genes and for RNA reads with biological variation compared to the reference.

A future exploration of the impact of reference bias on differential expression calls in wheat will be useful. Reference bias may have little impact on differential expression between conditions or across tissues within a single genotype, as, even if incorrectly quantified, the ratio of estimated expression between conditions/tissues should remain very similar regardless of reference. However, this needs to be assessed formally. If interested in homoeologue expression balance, however, unequal divergence of homoeologues relative to the reference will lead to incorrect findings. Reference bias also makes complex patterns more difficult to discern. For example, in a previous study [25], we demonstrated how the rhythmicity of *ELF3-1D* and *SIG3-1D* in a Cadenza timecourse RNA-Seq dataset was difficult to ascertain as the reads mapped so poorly to Chinese Spring. However, when using adding in the introgression to the reference, the reads mapped more correctly, and the rhythmicity could be accurately assessed.

Matching a sample to a more appropriate reference genome will become increasingly possible as genome assemblies for more wheat accessions become available. However, analyses involving two or more accessions require a common reference genome to which the RNA-Seq reads can be aligned. In this situation, or when the appropriate genome assembly is not available for within-accession analyses, it is important to exercise caution and check whether introgressed genes might be impacting conclusions drawn. In the long-term, it is important to work towards overcoming this issue of introgression-induced reference bias by implementing novel methodology.

### Using a pantranscriptome reference to reduce reference bias

Previous work has shown the benefit of using enhanced references or individualised references as targets for RNA-Seq mapping. [26] constructed an enhanced reference genome for human by including alternative allele segments at known polymorphic loci. Other publications have reported mapping to individualised genomes/transcriptomes by updating the reference with SNPs, INDELs and/or splice sites for each individual [27, 28]. Working with a multi-parent mouse population, Munger et al., 2014, found that mapping to individualised genomes increased the accuracy of eQTL assignment from 88.2% to 98.3% and corrected false-positive linkage signals. [29] constructed a pan-human consensus genome by calculating the consensus allele for each variant; this significantly improved the accuracy of RNA-Seq mapping when compared to the reference genome.

Our approach follows in this vein. However, individualised genomes or consensus genomes are not suitable for wheat as the degree of divergence introduced by introgressions prohibits the accurate genotyping necessary for creating said genomes. Instead, we built a pantranscriptome reference, that includes alternative transcripts from other wheat cultivars in the Chinese Spring reference transcriptome. The low resource requirements of kallisto regardless of reference size enables a highly scalable approach as more genome and transcriptome data are generated, while still running in a fraction of the time that alignment-based tools take to align to one reference genome.

The pantranscriptome reference corrects almost all expression values underestimated for genes belonging to an introgression present in the assembled pangenome cultivars and in a 1-to-1 relationship with a Chinese Spring gene. However, this approach does currently have limitations. The pantranscriptome reference will not currently contain all introgressions present across wheat accessions. As more genomes and/or transcriptomes are sequenced, transcripts can be added to the pantranscriptome reference to broaden the scope of genetic variation covered. This may lead to a saturation point at which most of the commonly segregating variation is captured within the reference and it can be considered complete for most use cases. This approach also only addresses errors caused by divergent genes and not those caused by copy number variation. Developing a way to overcome this limitation may prove very useful but is challenging because it requires resolving complex orthologue and paralogue relationships, and it is unclear how novel genes and genes with varying copy number between cultivars should be represented in the transcriptome reference.

Different solutions entirely to the problem of RNA-Seq reference bias in wheat may emerge as being superior. For example, the field of graph genomes is developing rapidly [30], including methods to align RNA-Seq reads to a graph genome [31]. However, graphs for genomes as large and as complex as wheat are yet to be created successfully. It is also a much heavier-weight solution compared to the pantranscriptome pseudoalignment approach. At the very least, our approach provides a temporary way to improve the accuracy of RNA-Seq alignment, particularly for those genes comprising the core genome. With further development and the incorporation of new data, it may evolve into an alternative, more lightweight approach to emerging graph-based methods.

### Examining reference bias in experimentally-generated RNA-Seq data

Using the valuable dataset generated by He et al. (2022), we were able to show that reference bias is present in experimentally-generated datasets as well as simulated datasets. The diverse nature of the wheat accessions sequenced may have made this work particularly prone to the effects of reference bias; after all, we demonstrated that divergent regions, likely introgressions, are abundant across the accessions. However, the ubiquity of introgressions is not exclusive to this set of accessions as introgressions are common across most wheat germplasm, including Elite cultivars. Indeed, wheat accessions containing diverse introgressions are very important in wheat research as it may be the source of beneficial variation for breeders, not to mention sources of insight into the evolution of wheat genomes.

The homoeologous sets of genes showing lack of expression correlation identified in He et al. (2022) were enriched in genomic regions identified as introgressed or deleted in many of the accessions with 78.2% falling in such regions. We also showed that most of these genes had much higher expression when using the pantranscriptome reference instead of the Chinese Spring reference. Using the pantranscriptome reference also increased the SCC scores calculated between homoeologue pairs. These findings may alter the interpretation of why these genes are associated with productivity traits. While some of these triads may still exhibit genuine dysregulation of homoeologues and homoeologue dosage effects, it is likely that, for at least some of these genes, variation in the gene sequence itself is underlying this trait variation, rather than alteration of expression dosage between homoeologues. This also has implications for the evolutionary and selection mechanisms implicated in controlling these traits.

To more precisely examine how the quantification of introgressed genes changes with the reference used, we focused on genes in the chr1D introgression due to its presence in around a quarter of the accessions and constant size across accessions possessing it. We showed that when using Chinese Spring as the reference, it appears as though introgressed genes are less expressed than the wheat orthologues they replaced. However, when using the pantranscriptome reference, which contains the introgressed gene models as the pangenome cultivar Jagger also contains this introgression, there is no significant difference between the expression of these genes. Correcting the quantification of these genes also altered the estimated expression balance across triads in which the D homoeologue is introgressed by raising the estimated expression of the D homoeologue. This has implications for any RNA-Seq studies using wheat accessions containing introgressions, and also more specifically for studies looking at the expression of introgressed genes and what mechanisms underlie the phenotype they confer.

## Methods

### Read simulation, alignment, and quantification

Reads were simulated from the longest transcript from each HC gene in Chinese Spring RefSeq v1.0 (with RefSeq v1.1 annotation) and the nine pseudomolecule genome assemblies [17] if the transcript >= 500bp. Wgsim from samtools v1.9 [32] was used to simulate 1000 pairs of 150bp reads per gene with an insert size of 400bp and no errors.

The kallisto index was produced from the CDS sequences from the RefSeq v1.1 high-confidence gene annotations using kallisto v0.44.0 [3]. Reads were pseudoaligned to this index using 100 bootstraps and default settings. Read counts and TPM values were summed across transcripts to generate gene level counts and TPM values.

To construct the pantranscriptome reference, we first ran Orthofinder [33] with standard parameters to define orthogroups based on the longest isoform protein sequences of the HC genes from Chinese Spring and the nine chromosome-level pangenome cultivars. If a gene was found in a 1-to-1 relationship with a Chinese Spring gene, its transcripts were added to the Chinese Spring RefSeq v1.1 HC transcript fasta file. A kallisto index was built and reads pseudoaligned as above. Read counts and TPMs were each summed across all transcripts of a gene and its 1-to-1 orthologues using the custom python script *sum_orthologue_transcript_counts.py* (supplementary data) to generate gene-level counts.

The STAR index was built for RefSeq v1.0 with the RefSeq v1.1 HC gene annotation using STAR v2.7.6a [2] using default parameters except for --limitGenomeGenerateRAM 200000000000 and --genomeSAindexNbases 12. The simulated reads from the 10 cultivars were aligned to this index using STAR and the predicted splice junctions from all were merged and then filtered to remove non-canonical junctions, junctions supported by 2 or fewer uniquely mapping reads and reads already annotated in the original genome annotation. The index was rebuilt using these discovered splice sites in addition to the annotated splice sites. The simulated reads from the 10 cultivars were aligned to this new index with parameters --quantMode TranscriptomeSAM and --outSAMunmapped Within. Gene-level read counts were generated using RSEM v1.2.28 [34].

For read count comparisons between self-mapping and cross-mapping, the following criteria were used to determine whether a gene was present in the analysis. For self-mapping, all genes from which reads were simulated were used. For cross-mapping, genes from which reads were simulated in that cultivar and that are in a 1-to-1 relationship with a gene in Chinese Spring from which reads were also simulated were used.

### Defining triad balance

Triads in Chinese Spring were taken from Ramirez-Gonzalez et al. (2018). For each cultivar, triads were retained if all three homoeologues were used to simulate RNA-Seq reads. Triad balance was computed in the same way as [6] except for the use of read counts rather than TPMs due to the way we simulated the reads. The relative read count of each homoeologue within a triad was calculated as follows:

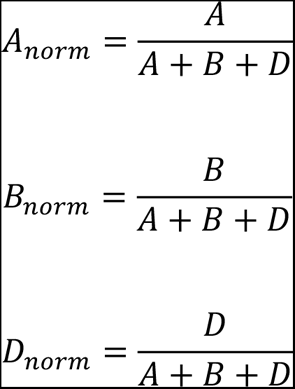

where A, B and D are the read counts of the A, B and D homoeologues, respectively. Euclidean distance was then used to calculate the distance between each set of normalised expression values across a triad to an ideal normalised read count bias for each of seven categories (Table 1). A triad is assigned to an expression bias category by selecting the category with the shortest Euclidean distance between the observed and the ideal bias.

**Table 1.** ideal normalised read count bias for each triad expression category:

| Category | A | B | D |
| --- | --- | --- | --- |
| Balanced | 0.33 | 0.33 | 0.33 |
| A suppressed | 0 | 0.5 | 0.5 |
| B suppressed | 0.5 | 0 | 0.5 |
| D suppressed | 0.5 | 0.5 | 0 |
| A dominant | 1 | 0 | 0 |
| B dominant | 0 | 1 | 0 |
| D dominant | 0 | 0 | 1 |

### Calculating CDS identity

Blastn from blast+ v2.7.1 [35] was used to align the nucleotide sequence of the longest transcripts of pairs of orthologues between Chinese Spring RefSeq v1.1 and Lancer. The identity of the best hit between pairs was taken and binned into 5Mbp genomic windows.

### Binning incorrectly quantified genes

The RefSeq v1.0 genome was split into 5Mbp genomic windows using bedtools makewindows [36] and for each window, a score was calculated based on the number of under (read count < 500) and overestimated (read count > 1500) genes within that window:

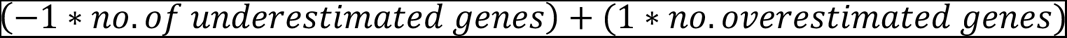

### Processing sequencing data from He et al., 2022

198 accessions had both leaf RNA-Seq data and enrichment capture short paired-end DNA reads. The RNA-Seq data from the 198 lines was pseudoaligned to both Chinese Spring RefSeq v1.1 and the pantranscriptome reference as above for the simulated reads. TPMs were summed across transcripts to generate gene level counts. Accessions GF25, GF270, GF32, GF37, GF41 and GF73 were excluded for RNA-Seq analyses as in He et al., 2022.

DNA reads were mapped to Chinese Spring RefSeq v1.0 (International Wheat Genome Sequencing Consortium (IWGSC) et al., 2018). The alignment was filtered using samtools (Li et al., 2009): supplementary alignments, improperly paired reads, and non-uniquely mapped reads (q <= 10) were removed. PCR duplicates were detected and removed using Picard’s MarkDuplicates (DePristo et al., 2011). Accessions GF294, GF342, GF366, GF380, GF381, GF383, GF38 were excluded for DNA analyses as in He et al., 2022.

### Using mapping coverage deviation to identify divergent regions of the genome

To generate sequencing reads for the pangenome cultivars, we simulated paired-end 150bp reads with 500bp insert and no errors from all fourteen genome assemblies (ArinaLrFor, Cadenza, Claire, Jagger, Julius, Lancer, Landmark, Mace, Norin61, Paragon, Robigus, Stanley, SY Mattis, and Weebil) generated by [15] to a depth of 10x using WGSim within samtools v1.9 [32]. Reads were mapped to RefSeq v1.0 as above.

The RefSeq v1.0 genome was split into 1Mbp genomic windows using bedtools makewindows [36]. Using the filtered read mappings for the pangenome cultivars and for the accessions from He et al. (2022), the number of reads mapping to each window was computed using hts-nim-tools [37]. To normalise by the sequencing depth of each line, read counts were divided by the number of mapped reads that passed the filters, producing normalised read counts c. Different windows of the genome have variable mapping coverage rates, so to compute coverage deviation we must compare each window to the same window in the other lines in the collection. Median normalised read counts, m, were produced, containing the median for each genomic window. Mapping coverage deviation was then defined for each line as:

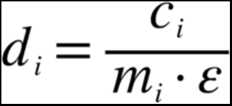

for window i ∈ {1, 2, …, n}, where ε is the median d value across the genome for the line. Statistically significant d values were calculated using the scores function from the R package ‘outliers’ with median absolute deviation (MAD) and probability of 0.99. Mapping coverage deviation and significance values were computed separately for the pangenome cultivars and for the accessions from He et al. (2022).

### Locating coordinates of introgression boundaries

To detect the precise locations of the chr1D, chr2A *Ae. ventricosa*, and the chr2D *Ae. markgrafii* introgressions in Jagger; and the chr2B *T. timopheevii* and the chr3D *Th. ponticum* introgression in Lancer, I used the alignments for the simulated Jagger and Lancer reads generated above. Read depths were binned into 5Mbp and 1Mbp windows using bedtools makewindows [36] and hts-nim-tools [37]. The window in which read depth drops, signifying the start/end of the introgression, was identified for each introgression and IGV was used to precisely identify the position where the coverage profile changes. To locate the location of the introgressions relative to the Jagger/Lancer genomes in order to identify which genes have been introgressed, I extracted Chinese Spring sequence 1Mbp either side of the precisely located border position (or until the end of the chromosome) for each introgression and aligned them to the Jagger or Lancer genome assembly using minimap2 [38] with parameters -x asm5. These alignments were used to determine the borders of the introgressed region as they appear in their donor genomes.

### Characterising the chr1D introgression donor species

Blastp from blast+ v2.7.1 [35] was used to align the *Ae. speltoides* proteins with the longest isoforms of the Jagger HC proteins. The best hit for each Jagger protein was kept. Paired-end Illumina DNA reads from *T. timopheevii* [39] were mapped to Chinese Spring RefSeq v1.0 using BWA mem v0.7.13 [40]. Samtools v1.4 [32] was used to filter the alignments to retain mapped reads, primary alignments, properly paired reads and uniquely mapping reads (q > 10). PCR duplicates were found and removed using the Picard Tools v2.1.1 MarkDuplicates function [41]. Read depths were binned into 5Mbp windows using bedtools makewindows [36] and hts-nim-tools [37] and divided by window length to account for windows at ends of chromosomes which are less than 5Mbp in length.

### Calculating SCC between homoeologues

SCC scores were calculated between AB, AD, and BD homoeologue pairs for triads where one homoeologue was in the set of genes showing lack of expression correlation identified by He et al. (2022). This was done using the cor.test function in R with the ‘spearman’ method and the lowest SCC value of the three comparisons was taken. Triads were excluded if any of the homoeologues were not found in the HC RefSeq v1.1 annotation.

## Declarations

### Ethics approval and consent to participate

Not applicable

### Consent for publication

Not applicable

### Availability of data and materials

The pantranscriptome reference, along with a python script to sum expression counts across all transcripts of a given Chinese Spring gene and its 1-to-1 orthologues, can be found in the Supplementary Data shared available at https://doi.org/10.6084/m9.figshare.24242767.

The RNA-Seq data and DNA sequencing data from He et al. (2022) reanalysed here are stored in the NCBI SRA under project codes PRJNA670223 and PRJNA787276.

### Authors’ contributions

BC conceived of the study and conducted analysis, prepared figures, and wrote the manuscript. TL identified 1-to-1 orthologues between Chinese Spring and the wheat pangenome cultivars. AH provided supervision and edited the manuscript. EA was involved in discussions and edited the manuscript. All authors read and approved the final version of the manuscript.

## Acknowledgements

We would like to thank Jose De Vega and Rachel Rusholme-Pilcher for providing feedback on an earlier version of the manuscript.

