## Supplementary Figures for "Introgressions lead to reference bias in wheat RNA-Seq analysis"

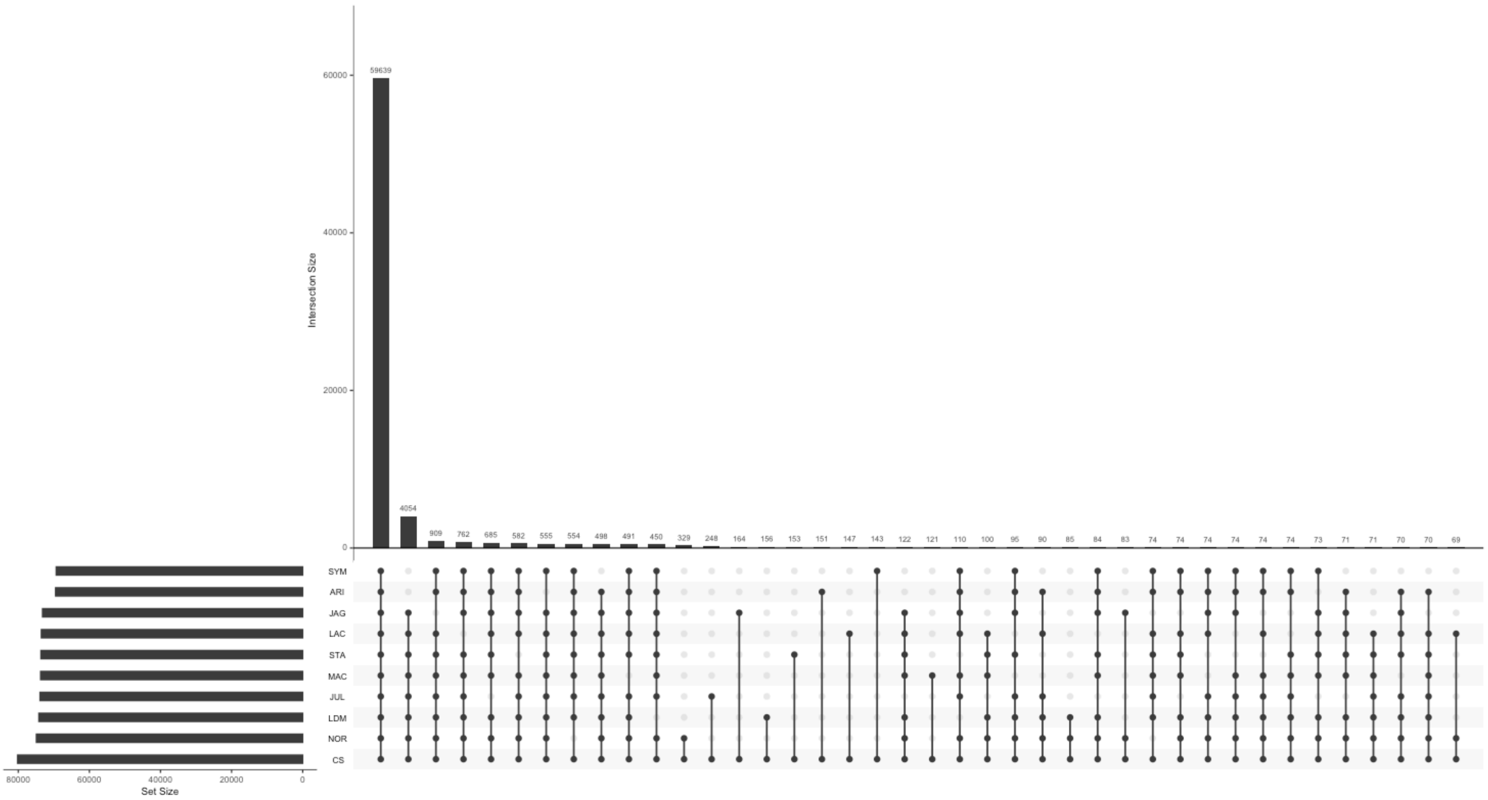


**Supplementary Figure 1. Upset plot of 1-to-1 orthologue assignments used for the construction of the pantranscriptome reference.**

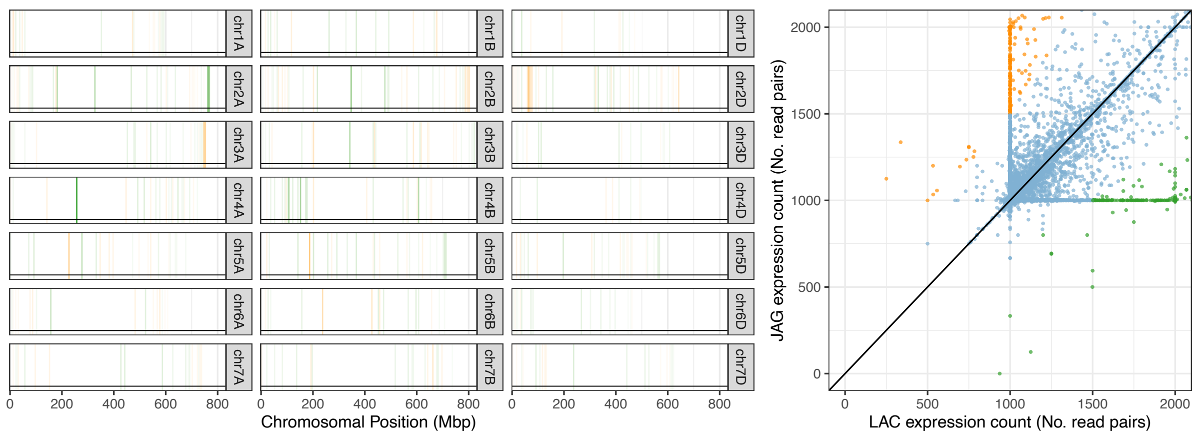


**Supplementary Figure 2. Remaining incorrectly quantified genes after correction using the pantranscriptome reference.** Scatter plot shows expression counts for simulated reads of Lancer-Jagger orthologue pairs when using kallisto with the pantranscriptome reference. Genes are considered incorrectly quantified if their estimated read count is 1.5x or 1/1.5x the other cultivar. The chromosome plot shows the distribution of incorrectly quantified genes in 5Mbp windows, coloured by the cultivar in which the estimated expression is lower; orange blocks are underestimated in Lancer compared to Jagger, while green blocks are underestimated in Jagger compared to Lancer.

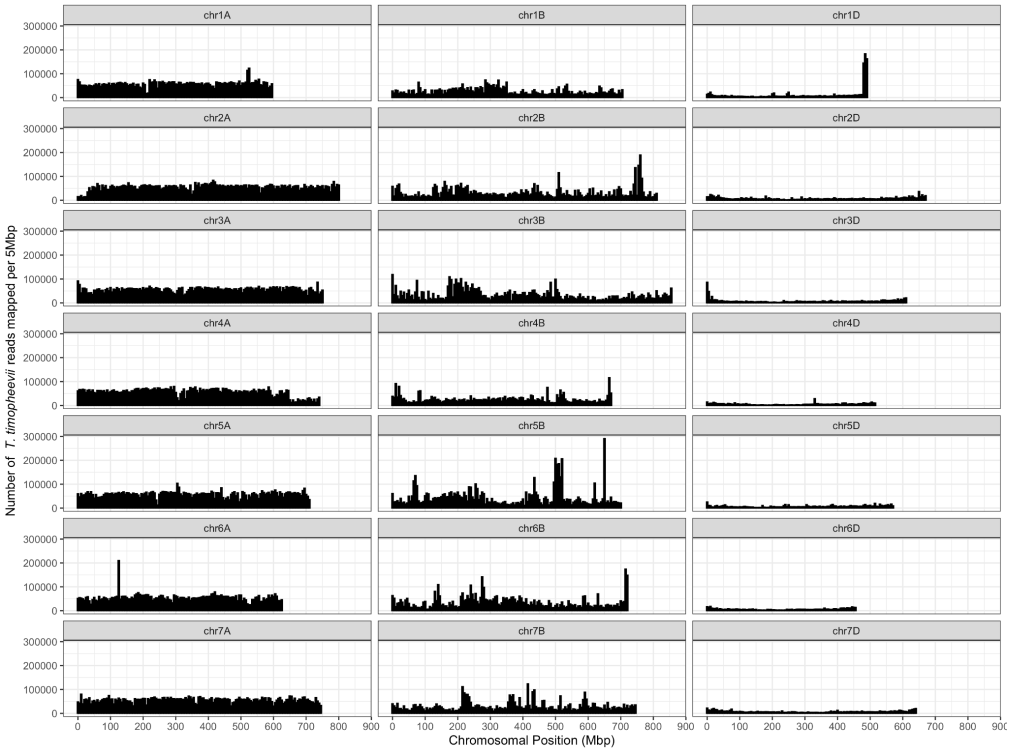


**Supplementary Figure 3. Reads from *T. timopheevii* accession P95 mapped to *T. aestivum* cv. Jagger and binned into 5Mbp genomic windows.** Number of mapped reads were divided by the length of window to accurately reflect read density at the final window of each chromosome. The chr1D introgression with a putative origin of *T. timopheevii* is at 481585620-493450010. *T. timopheevii* is a tetraploid with genomes related to the A and B subgenome of wheat. This is reflected in the greater mappability of *T. timopheevii* reads to the A and B subgenomes than the D subgenome. However, the higher read count across the 1D introgression suggests this region is more similar between *T. timopheevii* and the introgression than between its subgenomes and the A and B subgenomes, lending support to the donor of this introgression being *T. timopheevii*.
